# GIGANTEA gene expression influence leaf senescence in *Populus* in two different ways

**DOI:** 10.1101/2021.05.28.446188

**Authors:** Nazeer Fataftah, Pushan Bag, Domenique André, Jenna Lihavainen, Bo Zhang, Pär K Ingvarsson, Ove Nilsson, Stefan Jansson

**Affiliations:** Umeå Plant Science Centre, Department of Plant Physiology, Umeå University, Umeå, Sweden; Umeå Plant Science Centre, Department of Forest Genetics and Plant Physiology, Swedish University of Agricultural Sciences (SLU), Umeå, Sweden; Department of Plant Biology, Swedish University of Agricultural Sciences, Uppsala, Sweden

## Abstract

GIGANTEA (GI) genes have a central role in plant development and influence several processes such as light signaling, circadian rhythm and abiotic stress tolerance. Hybrid aspen T89 (*Populus tremula x tremuloides*) trees with low GI expression through RNAi have a severely compromised growth. In order to study the effect of reduced GI expression on leaf traits with special emphasis on leaf senescence, we grafted GI-RNAi scions onto wild type (WT) rootstocks and managed to restore scions’ growth. The RNAi line had distorted leaf shape and reduced photosynthesis, probably caused by modulating phloem or stomatal function, increased starch accumulation, higher carbon-to-nitrogen (C/N) ratio and a reduced capacity to withstand moderate light stress. GI-RNAi also induced senescence under long day (LD) and moderate light conditions. Furthermore, the GI-RNAi lines were affected in their capacity to respond to “autumn environmental cues” inducing senescence, a type of leaf senescence with characteristics different from senescence induced directly by stress under LD conditions. Whereas Overexpression of GI delayed senescence. The two different effects on leaf senescence were not affected by the expression of FT (Flowering locus T), were “local” – they followed the genotype of the branch independent on the position in the tree – and trees with modified gene expression grown in the field were affected in a similar way as under controlled conditions. Taken together, GI plays a central role to sense the environmental changes during autumn and determine the appropriate timing for leaf senescence in *Populus*.

**One sentence summary:** Leaf senescence is a complex process that is not well understood, but this paper shows that changing the expression of one gene could influence leaf senescence in *Populus* trees in two separate ways.

## Introduction

Every year deciduous trees go through a seasonal cycle of bud flush, growth, vegetative growth cessation, leaf senescence, dormancy, and cold hardiness. This cycle is important for surviving the boreal winter and has a big effect on the overall energy and nutrient balance. To properly respond to seasonal cues, trees have developed a complex regulatory network that integrates internal and external factors such as age, photoperiod, temperature, and nutrient status to set the appropriate time to switch from a stage of growth and photosynthesis to a stage of survival and nutrient remobilization. Obviously, there is a tradeoff between growth and survival/remobilization causing an evolutionary pressure to optimize the timing of phenological events and this has resulted in strong local adaptation of aspen (*Populus tremula*) phenology traits like bud set, which can also be observed in for example genome-wide association studies (GWAS) (Wang et al, 2018).

The regulation of autumn leaf senescence in trees is, in comparison to growth arrest and bud set, less well understood. At least in aspen, a tree in a given environment initiates the senescence process on almost the same date every year (e.g. Keskitalo et al, 2005) regardless of weather – i.e., it is under as strict seasonal control -however unlike for bud set photoperiod seems not to be the immediate, or only, trigger for senescence (Michelson et al. 2017). Temperature also affects autumn senescence in aspen (Fracheboud et al. 2009; Wu et al., 2018), but its effect is complex and probably less important than light for initiating autumn senescence under natural conditions. Accumulation of photosynthates (Lihavainen et al, 2020) and the level of different nitrogen species could also affect aspen leaf senescence, and severe drought or pathogen attack could of course initiate senescence at any time. Disentangling the different internal and external factors influencing senescence on the level of an individual tree has turned out to be a challenging task, not to mention understanding changes on landscape level (Zani et al., 2020). However, understanding the internal and external factors influencing autumn senescence is a key to predict how climate change will modulate senescence in different tree species, hence for modelling effects of climate change.

The ‘critical day length’ regulates different aspects of plant growth, e.g. flowering and vegetative growth (Andres and Coupland 2012; Pin and Nilsson 2012). The components of photo-periodically controlled flowering in annuals are well described, whereas knowledge of photoperiodic control in trees is more fragmented although the same components appear to be involved. Growth cessation and bud set in *Populus* is induced by the shortening photoperiod in the autumn and are dependent on light input via phytochrome (PHY) and the circadian clock components LHY1, LHY2 and TOC1 through the CO/FT module (Olsen et al. 1997; Böhlenius et al. 2006; Ibáñez et al. 2010; Ding et al., 2018; Miskolczi et al., 2019). GIGANTEA (GI) and GI-like (GIL) proteins control, in interaction with FKF1 and CDF1, seasonal growth cessation in *Populus* through regulation of the *FT2* gene (Ding et al. 2018). However, GI is involved in many other aspects of plant physiology; drought tolerance, miRNA processing, chlorophyll accumulation, cold tolerance, salt tolerance, herbicide resistance, phloem function and starch accumulation (Mishra and Panigrahi, 2015). Some of these effects may be consequences of the influence on circadian clock and phenology and separating primary from secondary effects is not an easy task. To study whether GI expression affects photosynthesis and leaf senescence in aspen, we set up a series of experiments using transgenic lines with modified expression of photoperiodic components in *Populus*, in particular grafted to each other. We found that GI is involved in modulation of leaf physiology (e.g. gas exchange and C/N ratio) and lowering of GI expression could induce leaf senescence by photooxidative stress under moderate light, but also that GI expression modulates senescence in response to short days and cold nights, obviously through a second pathway. We use these findings to draw conclusions on the interaction between physiological traits mediated by GI expression, and leaf senescence, and how senescence is modulated at the whole tree level.

## Results

### Poor growth of GI-RNAi trees could be rescued by grafting scions onto wild type (WT) rootstock

GI-RNAi lines have previously been shown to have severely reduced growth under climate chamber conditions (Ding et al. 2018). We selected one of the lines (8-2) previously studied and used this line for most of our studies, however, we also included a weaker RNAi line (line 1-1a) and a line overexpressing GI (GI-ox), see below. We grew the GI-RNAi line 8-2 trees in the field (in southern Sweden), under natural conditions and found huge (ca. 10-fold) differences in the growth (both stem height and diameter) between three-year old GI-RNAi and the wild type (WT) trees (Fig. 1A). The dramatic decrease in the growth and the early bud set of GI-RNAi, both under controlled conditions and in the field, complicates studies of the effect of *GI* expression on tree physiology. However, given that GI regulates phenology through FT2, which is a mobile signal in *Populus* (Miskolczi et al., 2019), we tested whether grafting GI-RNAi (line 8-2) scions onto a WT rootstock (Fig. 1B, “simple grafting”), which allows mobile signals such as FT to move from WT leaves in the rootstock to the apex, could rescue the effects on growth and bud set. Self-grafted WT trees were included as a control, and the grafted trees were grown under long day (18 h light; LD^18h^) conditions. Growth of the GI-RNAi scion was efficiently rescued by grafting onto WT rootstock, there was a dramatic difference between un-grafted and grafted GI-RNAi scions (Fig. 1C), even more obvious in the second or third growth cycle after bud set/dormancy break (Fig. S1A; Fig. 1E). The GI-RNAi scion expressed the *GI* gene to ca 25 % compared to the levels in WT leaves on the rootstocks, and the mRNA levels of *FT2*-gene was low in GI-RNAi scions (Fig. 1D). Hence, the rescue of growth is likely to be caused by the movement of FT and/or other mobile signals like gibberellins from the rootstock to the scion. Grafting made it possible to study the effect of GI expression on several aspects of development and physiology such as leaf morphology and senescence.

**Figure 1.**
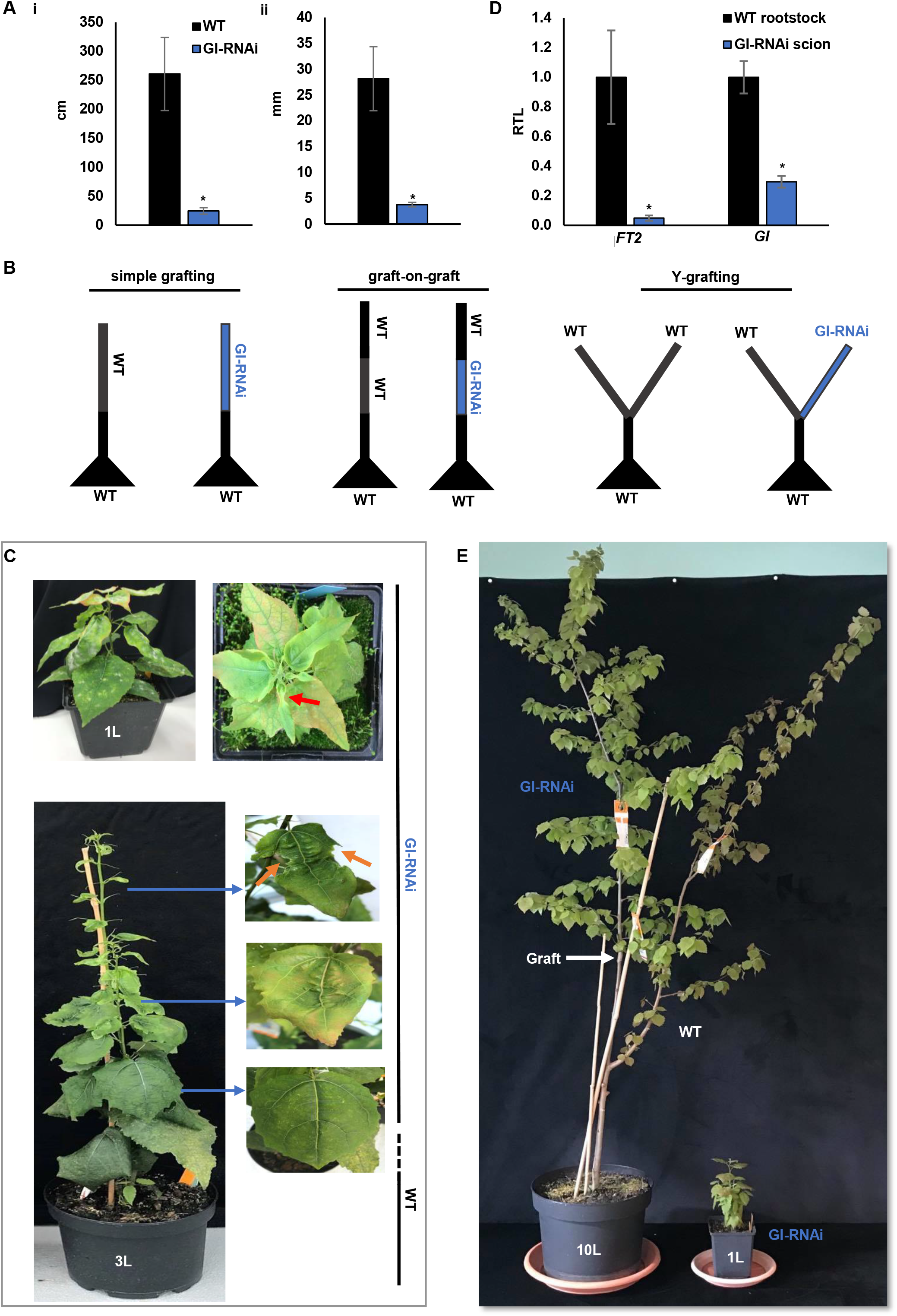
Rescuing the growth of GI-RNAi (line 8-2) by grafting on the WT rootstock. A, the growth of three years old GI-RNAi trees in the field; (i-ii) is the height and stem diameter, respectively; the bar is the average of 9 trees values ±SD. B, illustrations show the different grafting methods. C, upper pictures show ungrafted GI-RNAi phenotype; the red arrow points to the most affected area in the leaf; the lower pictures show the phenotype of grafted GI-RNAi scion on WT rootstock and the leaf shape in the right; the orange arrow points towards a necrotic part (black regions). D, the expression of *GI* and *FT2* in the GI-RNAi scions and WT rootstock of grafted trees under long day (LD^18^) conditions; RTL: relative transcription level; the bar is the average of 3 biological replicates ±SD. E, the grafted GI-RNAi scion on WT rootstock in the third growth cycle. The asterisk represents the significant difference using t-test; P <0.05.

While grafting could rescue the bud set and growth phenotype, leaf shape was still affected in GI-RNAi. The first leaves that were established in GI-RNAi after grafting had a normal shape but as the tree grew more leaves, they became increasingly aberrant with asymmetric growth around the main and secondary veins and leaves at the top of the shoot showed symptoms of necrosis/cell death (Fig. 1C). Accordingly, growing leaves of GI-RNAi (line 8-2) were less red, which also had lower anthocyanin and flavonol content than WT (Fig. S1A-D). Similar leaf shape patterns were also noted in the un-grafted GI-RNAi trees (Fig. 1C). In the second growth cycle, when the buds were flushed after dormancy, leaves emerging from the seasonal buds had normal shapes but as the shoot grew, subsequent leaves showed increasingly abnormal shapes indicating that the mobile signals from other tree parts affecting this phenotype were diluted as the tree grew.

### GI-RNAi leaves were sensitive to light and had lower CO_2_ assimilation rates

To study various aspects of leaf senescence in lines with modified GI expression we studied the leaf physiology of two RNAi and one overexpression line grown in various controlled environments. GI-RNAi (line 8-2) leaves were sensitive to the light when the trees were grown under LD^18h^ and standard light intensity in the greenhouse (200 µmol. m^-2^. s^-1^) causing chlorosis of GI-RNAi leaves (Fig. 2B), indicative of photooxidative stress. Also, the exposed leaves had more red colors (anthocyanins) between veins (Fig. 2B). However, unlike the growing leaf, in which anthocyanin is by default produced to prevent stresses, the anthocyanin of the RNAi mature leaf is a consequence of light stress. This photooxidative phenotype could be recovered (“regreening”) when the trees were moved to short day (14 h light; SD^14h^) and lower light intensity (150 µmol. m^-2^. s^-1^) conditions (Fig. 2A; B). In this context, GI has been proposed to influence stomatal conductance in *Arabidopsis* (Ando et al., 2013) and Edwards et al. (2010) showed that GI functions in veins influencing the wall ingrowth of phloem parenchyma cells. It is likely that functional synchronization of stomata and veins could cause the above-mentioned phenotypes of GI-RNAi leaves, e.g. green parts under LD^18h^ and asymmetry growth close to the veins (Fig. 1C; Fig. 2B). We shifted grafted trees in their second growth cycle from LD^18h^ to a growth chamber with SD^14h^ conditions with the intention to simulate autumn conditions (Fig. 2A). At multiple occasions we noted necrosis in GI-RNAi leaves already one week after the shift, while no such symptoms were found in the WT leaves (Fig. 2C). Necrosis was more pronounced around the main and secondary veins, suggesting a connection to phloem parenchyma cell malfunction in GI-RNAi (Fig. 2C). On the other hand, as the necrotic phenotype could possibly be associated with an increased air circulation – we noted that leaves facing the fan suffered more from necrosis than the less exposed leaves on the other side of the trees – it is also possible that stomatal malfunction may also contribute to this phenotype.

**Figure 2.**
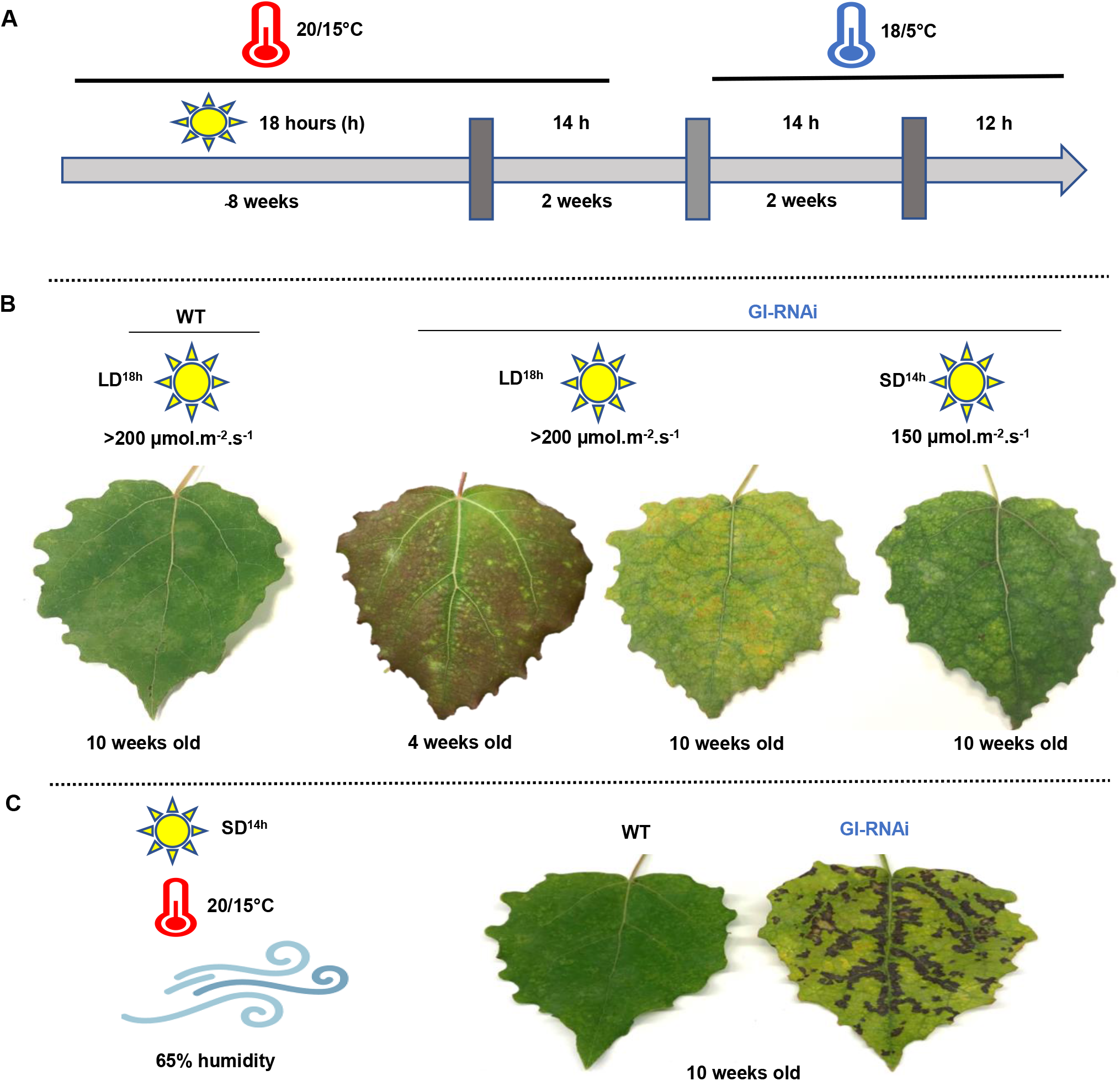
The leaves of grafted GI-RNAi (line 8-2) are sensitive to moderate light intensity. A, the indoor experimental setup simulating the autumn conditions. B, the leaf phenotypes of trees in their second growth cycle under differing photoperiod and light intensity; the right leaf of trees were transferred from LD^18h^ to SD^14h^ and lower light intensity; The upper bar represents the growth conditions. C, modulating the transpiration-caused necrosis/cell death in GI-RNAi leaves after one week under enhanced air circulation.

These observations indicated that photosynthetic capacity might be compromised in GI-RNAi leaves when incident light intensity exceeds the capacity to utilize the light energy under low photosynthesis causing photooxidative stress. However, the response was complex as there was spatial distribution between the regions close to the veins and interveinal regions (compare Fig. 1C and Fig. 2B; C). This prompted us to perform a detailed analysis of the photosynthetic performance of leaves of WT and GI-RNAi grafted plants. Trees were kept under LD^18h^ conditions and net CO_2_ assimilation rate (A_n_), stomatal conductance (g_s_), and the internal CO_2_ of the leaf (Ci) were measured weekly after the grafting using a LI-COR instrument. The same set of leaves – the first established after grafting -was measured throughout the experiment. Three weeks after grafting, WT and GI-RNAi leaves had relatively similar A_n_ (Fig. 3A). However, A_n_ decreased rapidly with time in GI-RNAi and after six weeks was only ca 20 % of WT (Fig. 3A). g_s_ was already lower in GI-RNAi three weeks after grafting and decreased even further with time (Fig. 3B) but Ci did not differ much either between the lines or over time (Fig. 3C). To further study the relationship between GI expression level and these photosynthetic parameters (A_n_, g_s_, and Ci), we also analysed a second GI-RNAi line (1-1a, grafted onto WT), where *GI* mRNA levels are only moderately decreased (Fig. S2), and a lines overexpressing *GI* (GI-ox, Ding et al. 2018). Line 1-1a showed little or no growth phenotype, no abnormal leaf shapes, and no signs of photooxidative stress under the conditions employed here, but they formed buds after two months already under LD^18h^ conditions. These lines did not differ from WT in gas exchange parameters (Fig. S3); obviously a more severe depletion of *GI* RNA was required to obviously compromise photosynthetic performance than to affect bud set.

**Figure 3.**
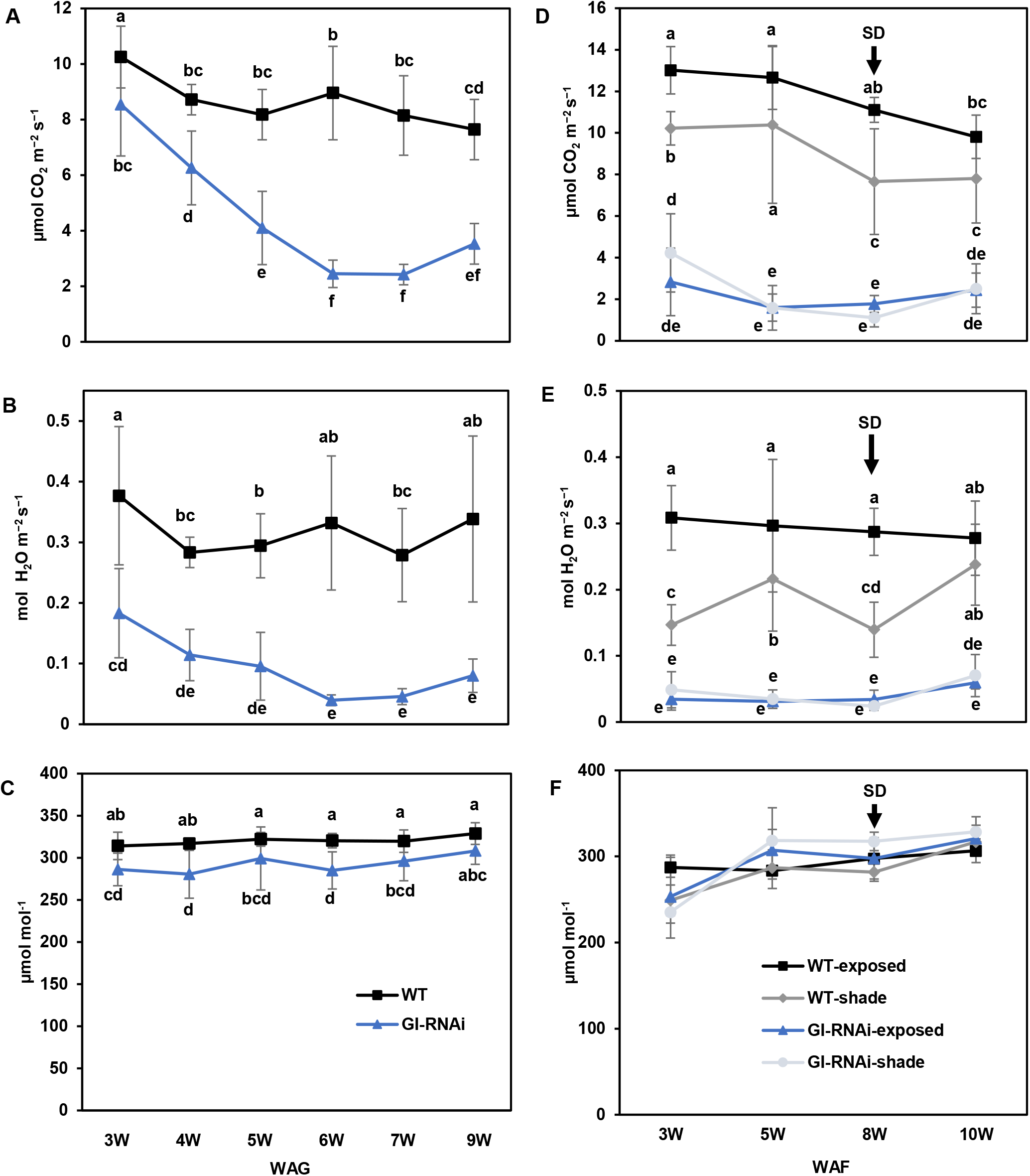
The gas exchange parameters of grafted WT and GI-RNAi (line 8-2) scions. A-C, the A_n_, g_s_, and Ci, respectively, of simple graft GI-RNAi or WT scions on WT rootstock under LD^18h^. D-F, the A_n_, g_s_, and Ci, respectively, of graft-on-graft trees that have WT scions on the top of grafted GI-RNAi; from eight weeks after flushing (WAF) the trees were subjected to SD^14h^. The data point is the average of 4-6 scion values ±SD. WAG: week after grafting. The different letters represent the significant differences; P <0.05.

### Intrinsic properties of GI-RNAi leaves do not depend on position in the tree

As GI modulates growth arrest, decreased CO_2_ assimilation in GI-RNAi (line 8-2) could potentially be the result of a changed sink demand. Also, to see if obvious mobile signals move upward from the RNAi parts, we grafted a WT scion on the top of grafted GI-RNAi, grafted on WT to create “graft-on-graft trees” (Fig. 1B). In following growth cycle after a short day/dormancy conditions, followed by bud flush under LD^18h^, trees continued growing like the control WT self-grafted trees. No difference in stem diameter of WT and GI-RNAi parts was observed (Fig. S4).

Here we also compared leaves that were shaded naturally by other leaves to study A_n_ and g_s_ of GI-RNAi under low light (when leaves were not obviously under light stress). Shading decreased the A_n_ and g_s_ of the WT leaves comparing to neighbouring fully exposed ones (Fig. 3D; E) but A_n_ and g_s_ were again much lower in both exposed and shaded GI-RNAi leaves, both under LD^18h^ or SD^14h^ conditions (see Fig. 2A for experimental setup), than in the WT leaves at the top of the same tree or self-grafted WT (Fig. 3 D; E; Fig. S5A; B). We also analysed starch accumulation, and GI-RNAi (line 8-2) leaves contained much higher starch levels than WT leaves (both below and above the GI-RNAi scion); this was evident both in growing and mature leaves (Fig. 4A; Fig. S4) particularly in exposed conditions; shaded leaves contained as expected less starch and the difference between GI-RNAi and WT was also less pronounced. The starch accumulation was also reflected in an increased fresh weight per area of the leaves by 7% (Fig. 4B) and their C/N ratio that was increased in exposed and shaded leaves but more pronounced in the exposed leaves of GI-RNAi leaves (Fig. 4C). It should also be pointed out that the starch accumulation of GI-RNAi leaves was less uniform than in WT, little starch accumulated along main and secondary veins, consistent with a role of GI in the function of stomata and/or veins (Fig. 4A). Taken together, GI-RNAi (line 8-2) leaves had lowered CO_2_ assimilation rate but higher C/N ratio and starch accumulation in exposed leaves also when growing between two sections of WT stems. Therefore, the effect is an intrinsic property of the GI-RNAi leaves, not of the sink/source activities. Furthermore, starch accumulation did not explain the lowering A_n_ and g_s_ in shaded leaves, indicating that stomatal, rather than phloem loading, malfunction was more important in explaining the lower A_n_.

**Figure 4.**
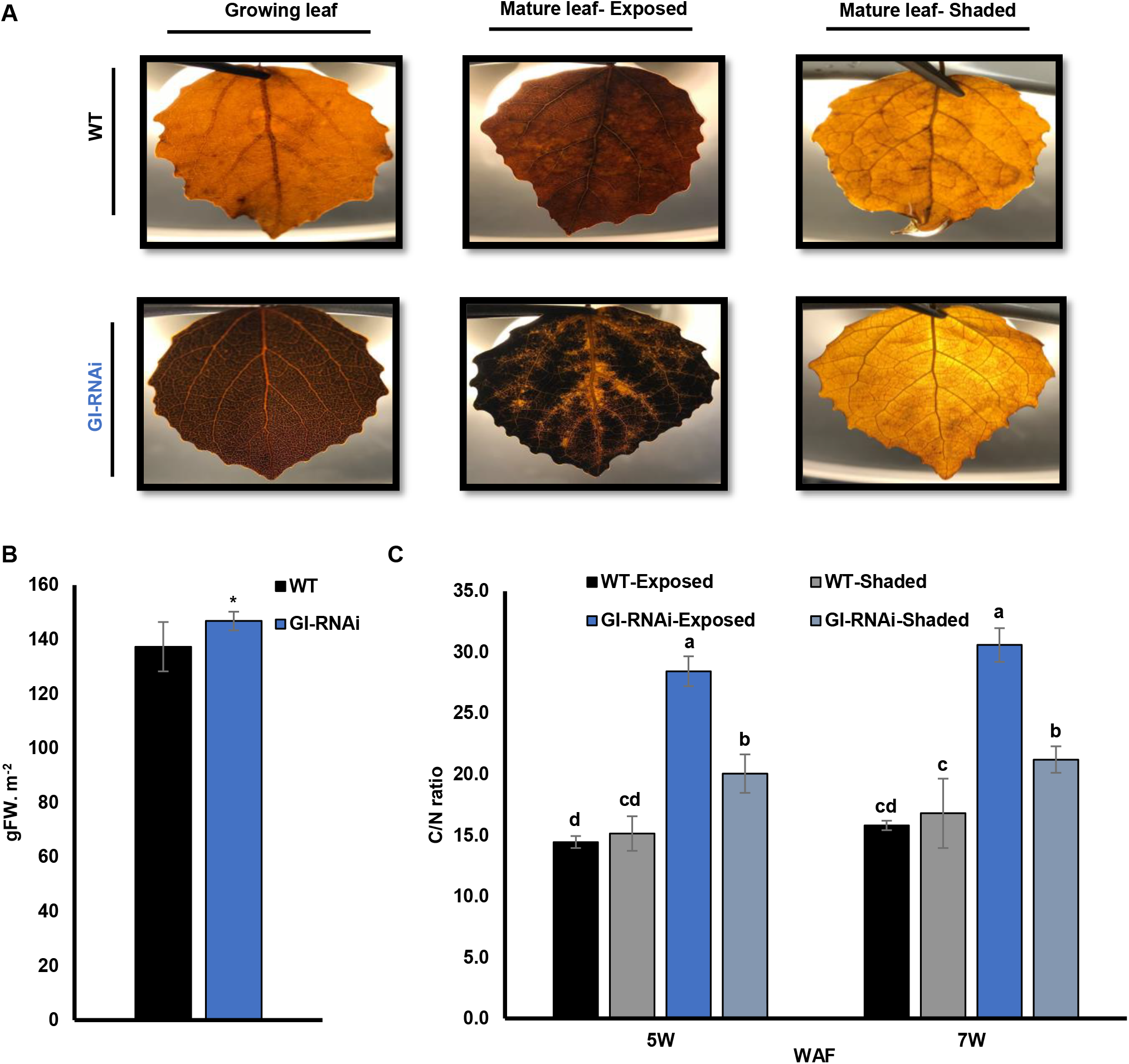
The starch levels and C/N ratio in WT and GI-RNAi (line 8-2) leaves on the same tree under LD^18h^. A, the starch staining in exposed and shaded leaves of the graft-on-graft trees; the pictures were taken with illumination from the rear for improved visualization. B, the fresh weight of leaves per area of the graft-on-graft trees; the bar is the average of 10 leaves ±SD. C, the C/N ratio in the leaves of the graft-on-graft trees; the bar is the average of 4 biological replicates from 4 trees values ±SD. The asterisk and different letters represent the significant differences using t-test or ANOVA analyses, respectively; P <0.05.

The chlorophyll and starch staining data indicated that there were spatial differences in chlorophyll fluorescence properties within GI-RNAi leaves. Analysing chlorophyll fluorescence using SPEEDZEN imaging system is a powerful way that can give information about different photosynthetic properties with spatial resolution, therefore we studied leaves of WT and GI-RNAi (line 8-2) exposed and shaded leaves with the age of two and seven weeks after flushing (Fig. 5) using imaging SPEEDZEN. In general, Fm in was, overall, lower in GI-RNAi leaves (Fig 5A) indicating that PSII was inhibited or quenched. Fv/Fm (Fig. 5B) tended also to be lower, indicating that PSII activity was somehow reduced as a consequence of quenching or photoinhibition. Some differences could also be noted in the amount of the fast (qE, Fig 5C) or slow (qZ/qI, Fig 5D) components of NPQ (Horton et al., 1996; Lambrev et al. 2012; Bag et al. 2020). The lower amplitude of NPQ (fast+slow) in GI-RNAi and their spatial distribution in the leaf could be caused by a reduced Calvin cycle activity, due to for example lowered activity of phloem transfer cells or stomatal malfunctions, Taken together, our data suggest that lowered GI expression primarily affects the photosynthetic dark reaction and that the effects on the light reaction is indirect.

**Figure 5.**
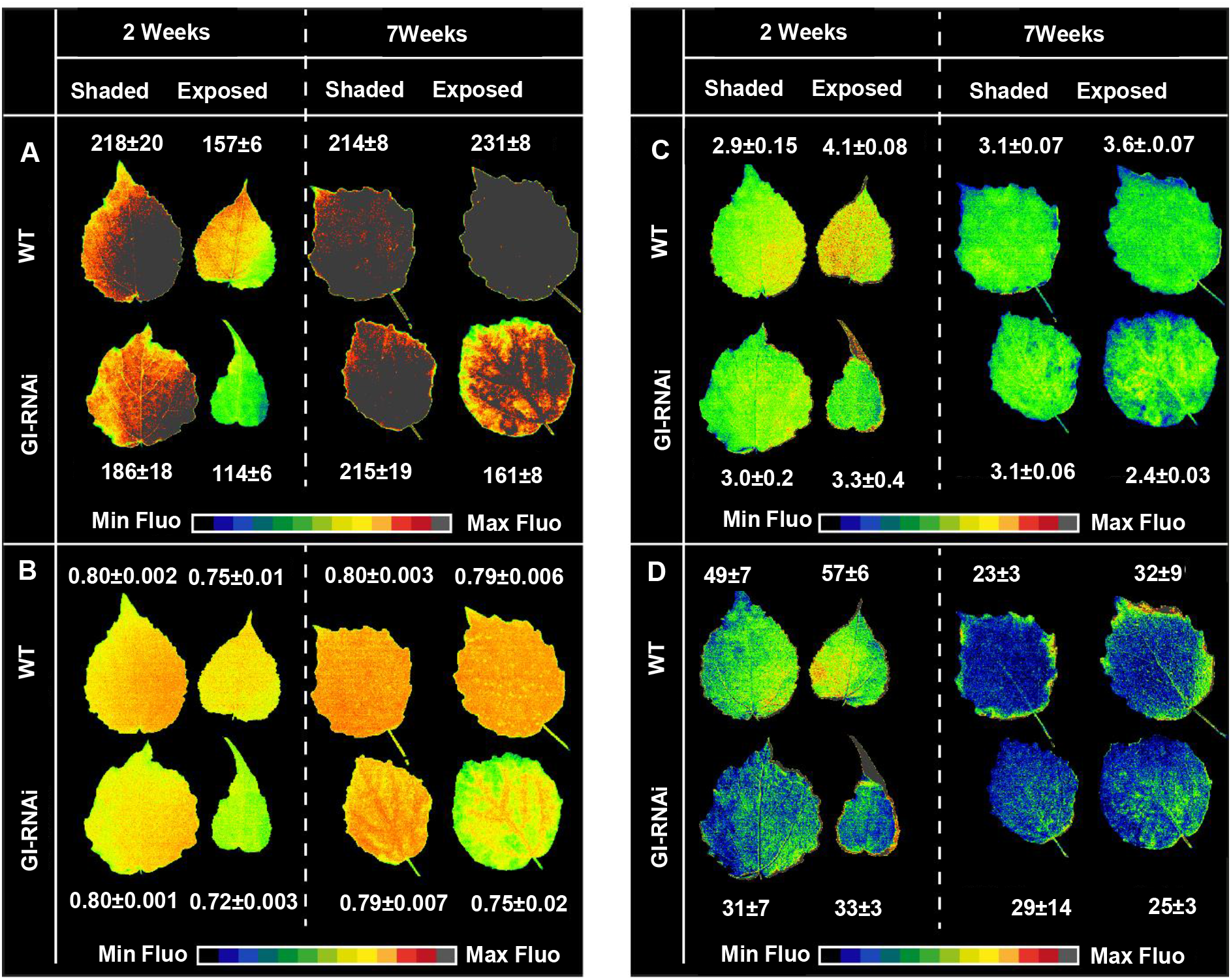
Photosynthetic performance of GI-RNAi (line 8-2) compared to wild type at two different stages (2 and 7 weeks) after bud flush of graft-on-graft trees for exposed or shaded leaves. A, maximum fluorescence (Fm, scale 1-200 r.u.) B, maximum quantum yield of PSII (Fv/Fm, Scale 0.01-0.80), and C, Fast component of non-photochemical energy dissipation (NPQ, scale 0.01-5) measured with saturating pulses (SP) under 2600 µmole constant actinic light. Representative images of fast and slow component of NPQ are shown from different SP applied during the measurement period. D, Slow component of NPQ measured from direct fluorescence from the last flash (14^th^) dark recovery period.

### GI-RNAi leaves affected leaf senescence by two different pathways

We have recently, using studies of the physiology and metabolism of senescing leaves of stem-girdled aspen trunks grown under natural conditions in the field along with non-girdled trunks on the same tree, been able to separate different patterns of leaf senescence in aspen. Girdling resulted in the early onset of senescence, overriding the “normal phenological control” of autumn senescence, i.e., that each tree genotype induces senescence at approximately the same date every year (Lihavainen et al. 2020). However, as we saw more similarities such as high C/N ratio and anthocyanin level between the stress-induced senescence in GI-RNAi (line 8-2) leaves growing in LD^18h^ and the senescence in girdled aspens, rather than in aspens undergoing typical autumn senescence in the field, we set out to induce autumn senescence in a growth chamber by changing light conditions and temperature. We included both grafted trees from their first and second growth cycle and the “graft-on-grafted” trees in this experiment. In addition, we included trees grafted with the weaker GI-RNAi line (1-1a), GI-ox and non-grafted trees of each genotype. To simulate autumn, the trees were moved from LD^18h^ to SD^14h^ and after two weeks night temperature was lowered (18/5°C day/night; T^18/5°C^) (the experimental setup is shown in Fig. 2A), resembling late August/early September conditions in Umeå.

Under these conditions, GI-RNAi (line 8-2) leaves were always senescent earlier than WT leaves, as was evident in exposed and shaded leaves and independent of the position in the tree (Fig. 6A). GI-RNAi leaves were typically shed two weeks before WT leaves. Furthermore, when ungrafted GI-RNAi line 1-1a trees were also subjected SD^14h^ T^18/5°C^, the leaves also became senescent much earlier than the WT (Fig. 6B), and the phenotype was reproduced when 1-1a was grafted as scion or rootstock with WT (Fig. 6C and Fig. S6A; B). Obviously, although this line grew and photosynthesized like WT (with no sign of photooxidative stress), senescence induced by SD and lowering temperature was affected, so this senescence trait – autumn senescence – was more sensitive to decreased expression of GI than the other type of senescence, which we hereafter will name “premature senescence”. On the other hand, GI-ox delayed the senescence by at least six weeks under SD^14h^ T^18/5°C^, both ungrafted and grafted branches displayed delayed senescence phenotype (Fig. 6B; C and Fig. S6C; D).

**Figure 6.**
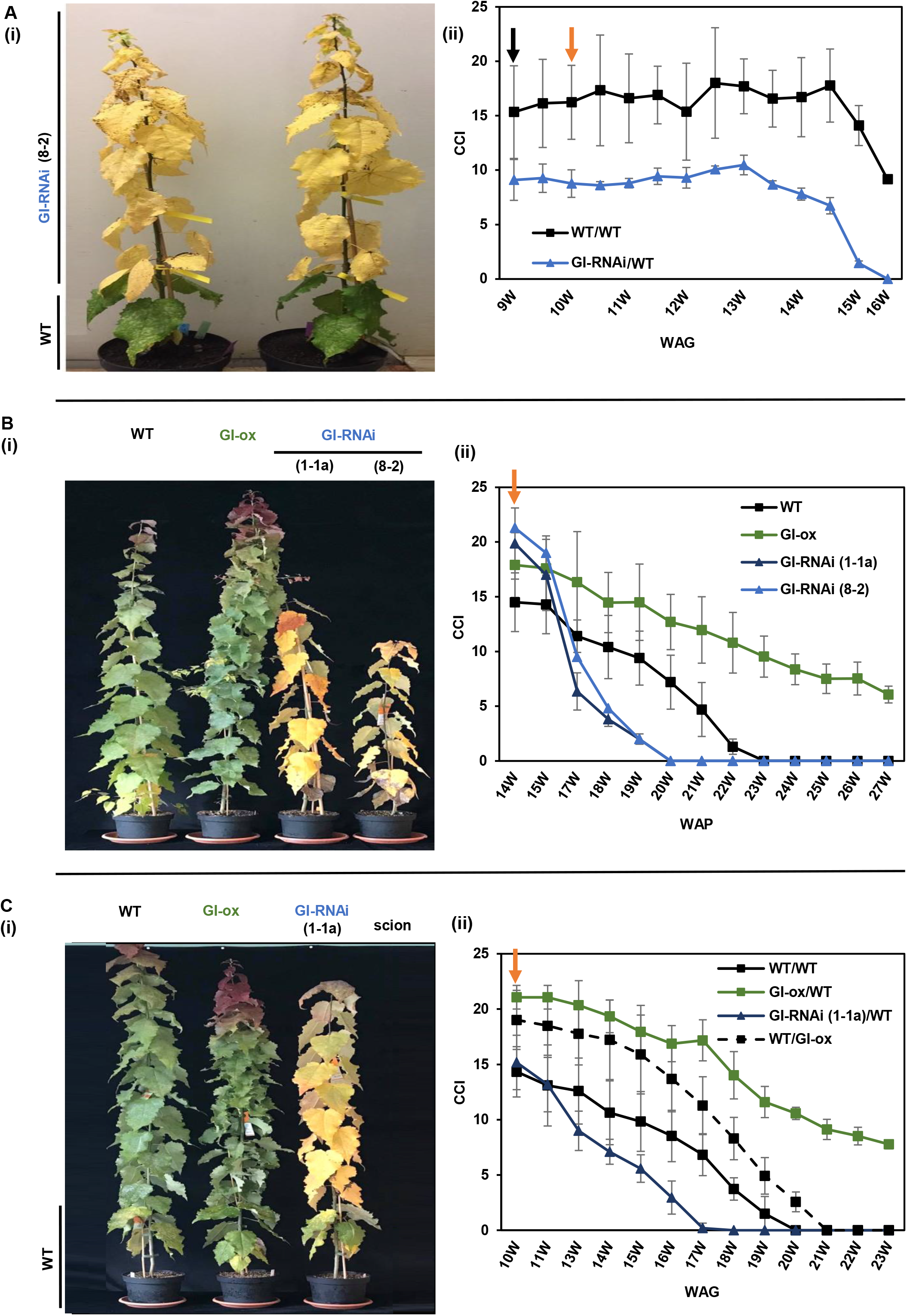
GI-RNAi leaves senescent earlier than the WT leaves under simulated autumn conditions. A, the senescence phenotype of grafted GI-RNAi (line 8-2) and WT scions on a WT rootstock under simulating autumn conditions; the photo was taken at fifteen weeks after grafting (WAG). B, the senescence phenotype of ungrafted WT, GI-ox, GI-RNAi (line 1-1a and 8-2); the right tree is only tree grew more than 15 cm from the line 8-2 trees; the photo was taken at twenty weeks after potting (WAP). C, the senescence phenotype of grafted WT, GI-ox, and GI-RNAi (line 1-1a) scions on a WT rootstock; the photo was taken at sixteen WAG. (B-ii) the CCI of ungrafted trees; (A, C-ii) the CCI of scions of grafted trees represented by corresponding left picture. The data point is the average of 4-6 trees values ±SD. The black arrow represents when the trees subjected to short days, and the orange arrow represents when the trees were subjected to cold nights.

We also observed that GI-RNAi (line 8-2) leaves, that senesced earlier than WT, seemed to indirectly affect the senescence behaviour of WT leaves; WT leaves that share the same tree with GI-RNAi leaves in the graft-on-graft plants had higher pre-senescence chlorophyll levels and entered senescence later than control grafted WT trees (Fig. 7). We believe this to be a consequence of improved nutrient status; as GI-RNAi leaves accumulate less nitrogen (and chlorophyll; Fig. 4C and Fig. 7), more nitrogen is available to WT leaves on the same tree during growth and even more so when the GI-RNAi leaves have started to senescence, and the remobilized mineral nutrients become available for the rest of the plant.

**Figure 7.**
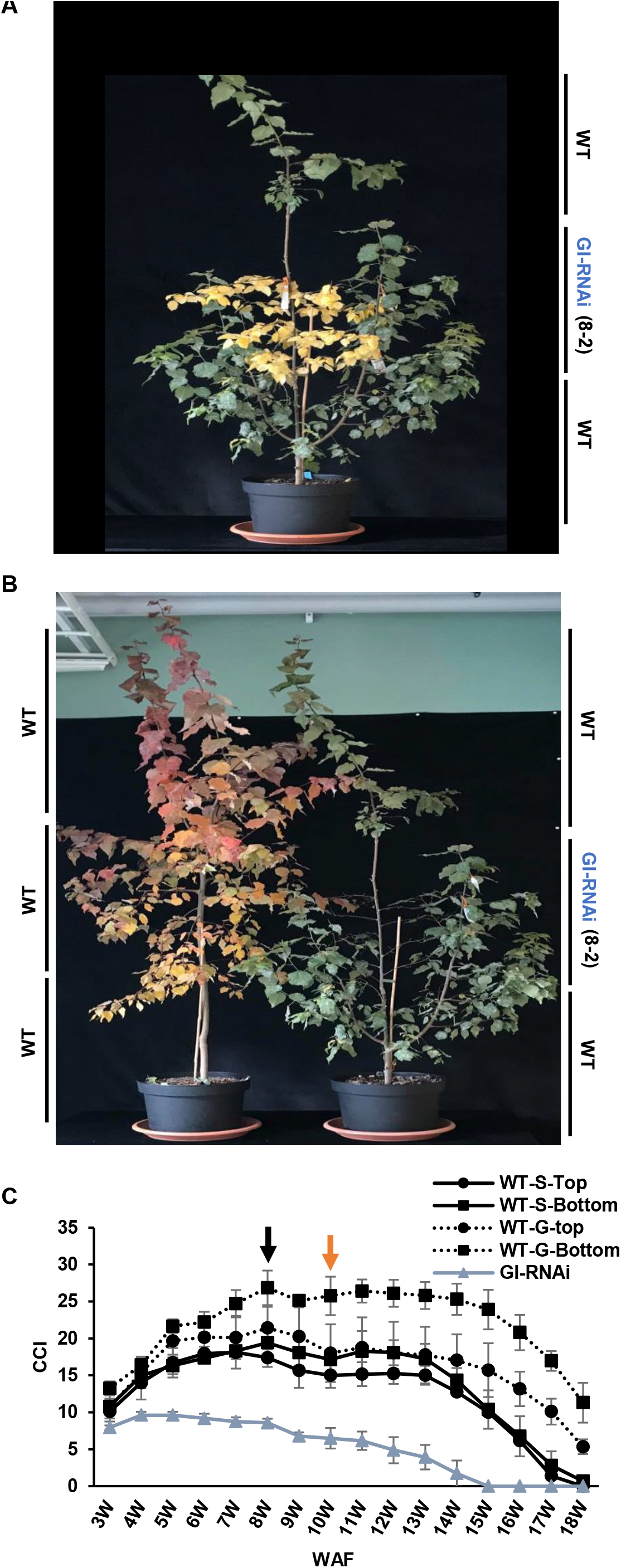
The senescence phenotype of graft-on-graft trees under simulated autumn conditions. A and B, the pictures represent the senescence phenotype in WT and GI-RNAi (line 8-2); the photos were taken at fourteen and sixteen weeks after bud flushing (WAF), respectively. C, the CCI of leaves graft-on-graft trees; WT-S: Self grafted WT. WT-G: WT scion grafted on GI-RNAi. The black arrow represents when the trees subjected to short day, and the orange arrow represents when the trees were subjected to the cold night. The trees were grown in 10 litre pots. The bar is the average of 4 trees values ±SD.

During the experiments when senescence was induced by SD and lowered temperature (SD^14-12h^ T^18/5°C^) – simulating autumn senescence – we noted differences in the patterns of leaf senescence in GI-RNAi leaves (in the stronger mutant lines; 8-2) compared to when senescence was induced under LD^18h^, “premature senescence”. “Autumn senescence” of GI-RNAi was rather uniform but started along the main and secondary veins in both exposed and shaded leaves than interveinal regions, while in some cases the veins stayed green similar to intervein than vein’s close regions. These regions were associated with the previously mentioned phenotypes such as reduced starch accumulation or photooxidative stress in the exposed leaves (Fig. 2; Fig. 4A). WT and line 1-1a senescence under “simulated autumn conditions” was more uniform. In LD^18h^, “premature senescence” in GI-RNAi line 8-2 started in intervein regions mainly in the exposed leaves. Under LD^18h^ conditions, leaves also produced more anthocyanin and flavonol compounds (Fig. 8B; C). The differences were obvious not only in the coloration of the leaves when visually compared but also when the chlorophyll fluorescence patterns were compared (Fig. 8D). Taken together, these observations show that a reduced expression of GI in *Populus* leaves affected two different pathways leading to leaf senescence, one that made the strong RNAi line (8-2) leaves more vulnerable to (photooxidative) stress already in the stage of active growth of the tree and associated with starch accumulation in the intervein regions. The second pathway was induced by shortened photoperiod and lowering the temperature, i.e., resembling conditions that induce the autumnal senescence that we study in the field. We also subjected WT and GI-RNAi (line 8-2) leaves, grafted onto a WT rootstock, to a SPEEDZEN time-course analysis over 15-weeks (LD^18h^ to SD^14h^-to-SD^14h^ T^18/5°C^). The results (Fig. S7) were consistent with lowering of GI expression both reduced the leaf’s ability to withstand photooxidative stress and changed their senescence behaviour when days became shorter and nights colder.

**Figure 8.**
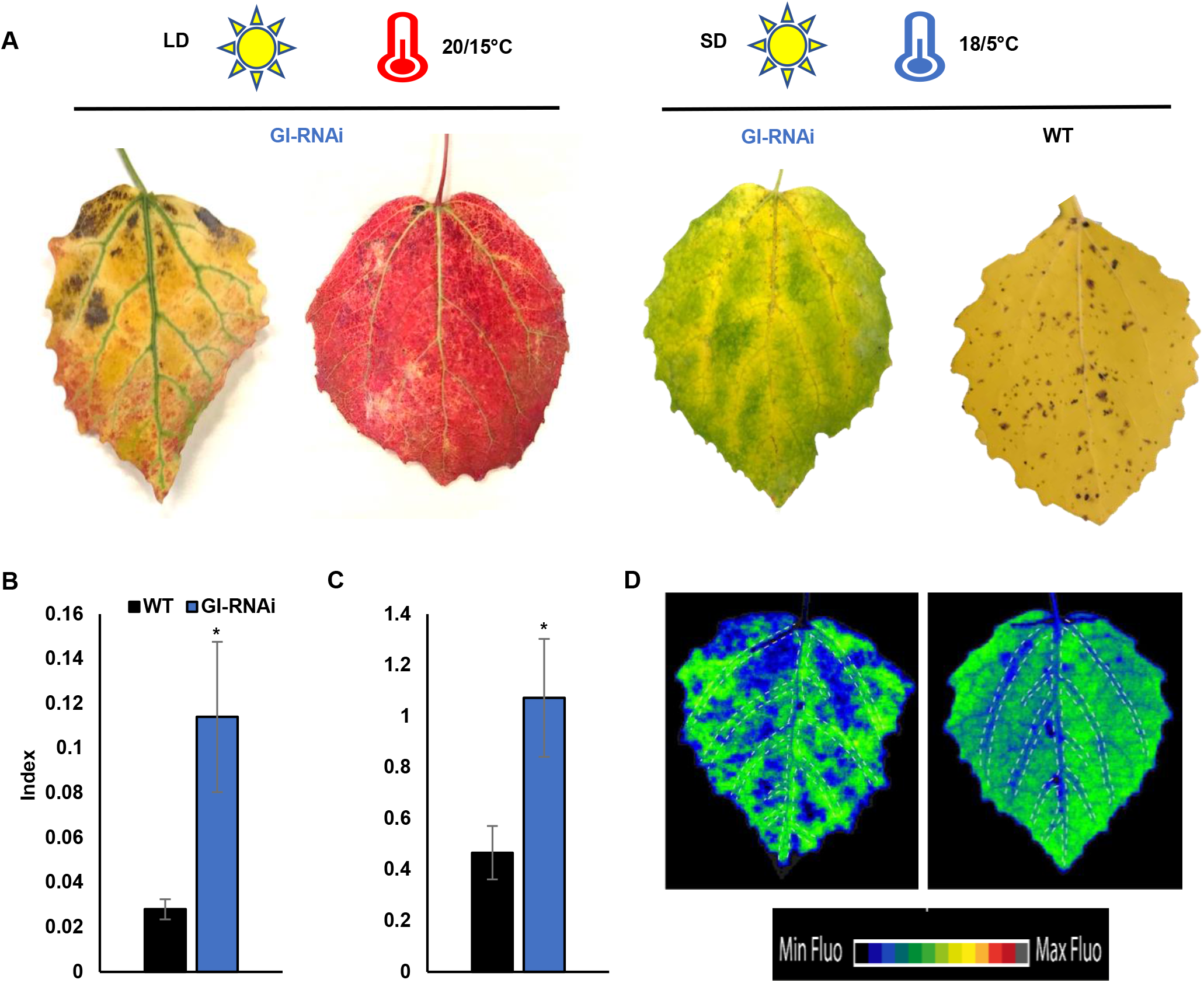
The two types of senescence in GI-RNAi (line 8-2) depending on the conditions. A, leaf senescence phenotype of GI-RNAi under LD^18h^ or short days and cold nights; WT leaf represents the typical uniform leaf senescence in the field and under simulated autumn conditions; the upper bar represents the growth conditions. B, the anthocyanin index of the leaves under LD^18h^ conditions; C, the flavonoid index of the leaves under LD^18h^ conditions. D, chlorophyll fluorescence spatial patterns in GI-RNAi leaves under LD^18h^ (left leaf) and short days and cold nights (right leaf); the dashed lines show the veins. The values are average of four biological replicates. The asterisk represents the significant difference using t-test.

### GI expression influence senescence also under field conditions, whereas FT expression did not

In the laboratory, under LD^18h^ conditions, GI-RNAi trees were able to grow if provided with a mobile signal from a WT rootstock (i.e grafted on WT) but were still affected in the progression of leaf senescence. To see if the same was true under field conditions we set up an outdoor experiment where trees were grown in pots but exposed to natural changes in light and temperature (in Umeå). Several different trees were used (in several replicates), a) GI-RNAi (line 8-2) grafted to WT and b) control grafted WT, both in their second growth cycle. For these trees WT buds of the rootstock were removed, producing trees with only one type of leaf after bud flush to check the senescence phenotype when the tree has only GI-RNAi leaves. In addition, we included trees with GI-RNAi and WT scions on the same WT rootstock, resulting in a “Y grafting” (Fig. 1B; Fig. S8B). In these trees, the WT scion in the Y graft grew faster and growth of GI-RNAi branches were supressed already under LD conditions, in contrast to the situation in simple grafted trees. Obviously, the WT scion was able to impose apical dominance over the GI-RNAi scion. After a cycle of dormancy/dormancy break, all trees were flushed in LD^18h^ T^20/15°C^ and moved to the outdoor in July (when the photoperiod was ca 20 hours). The GI-RNAi parts of all trees set bud already in July, while the WT in the two groups including the trees that had GI-RNAi scions did not form buds until October. GI-RNAi leaves senescent and were shed earlier than WT leaves (Fig. S8C; D) and under these conditions senescence also started around the veins of GI-RNAi leaves in September (Fig. S8C). The effect of GI on aspen leaf senescence under field conditions appeared not to be mediated through FT. When FT-RNAi, WT, and FT-ox (overexpressing FT) aspen trees were grown under outdoor conditions, FT-RNAi set bud more than one month before the other lines, but senescence was not affected (Fig. S9A). This was also confirmed by measurements of senescence in two-year old trees of FT-RNAi, WT, and FT-ox in a field experiment in southern Sweden (Fig. S9B), also in this case bud set was affected in FT-RNAi but senescence was not different from WT.

## Discussion

We still only have a shallow understanding of leaf senescence. Although it is a strictly regulated developmental program, much effort has been expended to identify genes regulating leaf senescence, how environmental conditions trigger the process, and which hormones are involved. Moreover, there isn’t much consensus on which genes, metabolites or pathways that provides the keys to the process. One obvious possibility is that if senescence is the default pathway for leaf development, which has to be prevented from being executed by a set of “blocking factors”, those that are removed could vary between conditions and species. Comparing genetically identical plant individuals experiencing different conditions is a useful way to understand complex phenomena, and has also been widely used to understand senescence, although alternative approaches could give additional information. We have, for example, recently shown that different parts of the same tree could induce senescence in slightly different ways. Girdling one of the trunks in a multi-stem aspen individual introduced changes in leaf pigment content, metabolites and senescence behaviour, in other words, this treatment induced variation in different parts of one single individual. Here we explore a different strategy by studying senescence process within a tree, in which parts of the tree have different genetic backgrounds exposed to the same environmental conditions. With this setup, we disentangle two modes of leaf senescence, expressed within the same tree. Reducing the expression of GI in controlled conditions was moderately stressful under moderate light intensity for the leaves and resulted in an earlier senescence and anthocyanin accumulation, presumably because of fundamental changes to leaf physiology and photosynthetic characteristics. In *Arabidopsis*, lowered GI expression leads to malfunction in phloem loading (Edwards et al. 2010), and our observations are consistent with a compromised phloem export also in *Populus*. Therefore, the early senescence we observed already under LD conditions in GI-RNAi (the stronger line; 8-2) could be analogous to the early senescence that can be induced by disrupted transport of photosynthates due to stem girdling (Lihavainen et al. 2020); lowered GI expression could disrupt transport out of the leaf. Another type of leaf senescence was observed when we exposed the trees to conditions that simulated autumn conditions: GI-RNAi leaves senescent earlier than WT leaves, but here the senescence process resembled more the “typical autumn senescence” that every autumn turns boreal forests colourful in a very consistent fashion. However, low GI expression also influenced the way how the leaf experienced “autumn conditions” that initiate leaf senescence in a manner that we have studied extensively over the years in aspen (Bhalerao et al., 2003; Andersson et al., 2004; Keskitalo et al., 2005; Fracheboud et al., 2009; Michelson et al., 2017; Edlund et al., 2017); a process that under natural conditions is initiated more or less the same date each year in a given genotype and largely unaffected by weather. This type of senescence is quite different on a macroscopic level: leaves senesce in a more uniform way in exposed and shaded leaves, and with less obvious signs of photooxidative damages. We believe that these two modes of leaf senescence have a wide physiological relevance but under most conditions are hard to study since they could occur in parallel, and our experimental manipulations – affecting the expression of GI or girdling experiments – just has made it possible to better distinguish between the two modes. A better distinction between different aspects of leaf senescence would be useful for the community, and the mere fact that we are uncertain how to name them illustrates the lack of a conceptual framework. We decided to use the term “premature senescence” although “stress-induced senescence” could be an alternative name.

We also use our results to draw other conclusions about autumn senescence. First and foremost, although premature/stress-induced senescence clearly could be local, it has previously not been clear whether autumnal senescence in a mature tree is local or systemic. Early observations that street lights may delay leaf fall (Matzke, 1936) have led to speculations that illumination of one part of the crown of a tree could keep that part green. The fact that this has not been possible to replicate by others (e.g. Sarala et al, 2013) or us (unpublished results) could be interpreted as if there is a systemic “senescence signal” that coordinates autumn senescence within a tree. However, the non-uniform senescence behaviour in our grafted trees showed unequivocally that, at least in *Populus*, senescence timing was strictly dependent on the genetic background of the branch. Secondly, the effect of GI on autumn-induced senescence is, in contrast to the effect on bud set, not mediated by FT whose expression level did not influence autumnal senescence, not even under field conditions. FT is probably indirectly involved, as growth cessation and/or bud set somehow predisposes the tree for sensing the illusive “autumn signal”. This is consistent with our previous findings that the light signal that is triggering induction of autumnal senescence in aspen is not daylength *per se* (Michelson et al. 2017), believed to act through FT (Ding et al. 2018). Instead, another light signal seems to be sensed and transduced through GI to trigger autumnal leaf senescence.

Our results do, however, also give information of leaf senescence not directly triggered by an “autumn signal”. Reducing GI expression resulted both in reduced growth, aberrant leaf development, and made the leaves sensitive to photooxidative stress. Grafting onto a WT rootstock that can provide the mutant with mobile signals such as FT rescued growth but not increased starch accumulation, increased C/N ratio of the leaves and senescence starting in intervein regions in the leaf under LD^18h^ (Fig. 4; 8). Lowering GI expression in *Arabidopsis* phloem leads to phloem transfer cells or stomatal malfunctions (Ando et al., 2013; Edwards et al. 2010), and our data suggest that the same is also happening in *Populus*, causing a reduced photosynthesis in GI-RNAi leaves and leading to the senescence phenotype. In general, a C/N imbalance is often associated with leaf senescence in (Wingler et al., 2006; Aoyama et al., 2014; Chen et al., 2015; Fataftah et al., 2018) but it is unclear whether the C/N imbalance that we noted in the leaves is a cause or the consequence of the changed photosynthetic performance. Stomatal malfunction may also cause an array of the effects, and photosynthesis gas exchange could be compromised, which would hamper photosynthesis, lead to downregulation photoinhibition and so on. The relation between GI expression and stress is, however, complex. For example, a knockout *gi* mutant is more resistant to herbicides, salt, or external H_2_O_2_ (Cha et al., 2019; Kim et al., 2013; Cao et al., 2006) and down-regulation of *GI*-like genes confers salt stress tolerance in *P. alba* x *glandulosa* (Ke et al., 2017). However, *GI* mRNA levels increase severalfold in cold treated *Arabidopsis* plants (Fowler and Thomashow, 2002), and the knockout mutant is more sensitive to low temperature (Cao et al., 2005). More studies are needed to find out how exactly GI expression relates to stress in the different plant species, developmental stage, light intensity, and whether this response is dependent-or independent-of stomatal conductance and carbohydrate status.

If not FT, what are the molecular players downstream of GI that regulate autumn leaf senescence? Furthermore, why do the regions close to veins sometimes senesce earlier, sometimes later in GI-RNAi leaves? Several reports suggest that the photoperiodic components are mainly expressed in stomata and vascular tissues (Imaizumi et al., 2005; An et al., 2004; Para et al., 2007; Adrian et al., 2010; Edwards et al., 2010; Kinoshita et al., 2011; Ando et al., 2013). Controlling nutrient and signal flow by stomatal function and/or vein function (including xylem unloading or phloem loading) could potentially explain how the light signalling/circadian clock controls leaf physiology. One potential connection between these pathways could be ethylene or ROS. Haydon et al. (2017) demonstrated that ethylene shortens the circadian period in *Arabidopsis*, conditional on the effects of sucrose and requiring GI. In addition, phytochrome interacting factors (PIFs), targeted by the GI pathway (Nohales et al., 2019), are involved in both ethylene biosynthesis and signaling pathway during dark induced leaf senescence (Sakuraba et al., 2014; Song et al., 2014).

To conclude, a reduced expression of GI in *Populus* leaves had dramatic changes in leaf physiology causing changes in for example leaf shape, C/N ratio, photosynthesis, and, most notably, leaf senescence through two different pathways, that appears to be independent of the expression of FT and bud set.

## Material and methods

### Plant Material and Growth Conditions

Hybrid aspen *Populus tremula L*. X *Populus tremuloides* Michx, clone T89 (WT), FT-RNAi, GI-RNAi lines (8-2 and 1-1a), and GI-ox that were characterized in Böhlenius et al. (2006) and Ding et al. (2018) were obtained from the tissue culture facility (Umeå Plant Science Centre) and potted in soil in three litre pots. The saplings were grown in the greenhouse with long day conditions (18/6 h day/night), temperature (20/15 °C day/night), light intensity (200 µmol. m^-2^. s^-1^) and 60% relative air humidity. The trees were fertilized weekly with 100 ml of diluted fertilizers (1:100; V:V) (SW Horto company).

### Grafting Experiments

Soil-grown plants were grown in the greenhouse, and then scions were grafted onto root stocks: the graft was covered by a plastic bag until the graft established. Different tree development stages of five, ten, and fifteen leaves were also tested. The grafted trees were initially grown under LD condition before transfer to different growth conditions.

### Chlorophyll, anthocyanin, and flavonol indices

Chlorophyll content index (CCI) of leaves (mean of five leaves) was determined for at least four independent trees from each genotype with a chlorophyll meter (CCM 200 plus, Opti-Sciences). The anthocyanin and flavonol indices were measured using Dualex Scientific+ (Force-A). For every biological replicate five leaves were measured and averaged.

### Simulating autumn conditions

The ungrafted and grafted trees were transferred from LD to SD (14/10 h; 20/15°C day/night) (Fig. 2A). After two weeks, the trees were subjected to cold night conditions (18/5 °C day/night), then after a further two weeks the photoperiod decreased to 12/12 h (day/night). The fertilization was stopped after three weeks in SD condition coinciding with the decrease in growth and the starting of dormancy stage.

### Second growth cycle

The trees that passed the simulation of autumn conditions were subjected to 5 °C temperature and photoperiod (8/16 h; day/night) for two months to break the dormancy. The trees were re-potted in ten litres pot and flushed again under LD conditions. A similar experiment setup that mentioned in the previous section was used to study leaf senescence under simulating autumn conditions for these trees.

### Starch staining and leaf weight measurement

For starch staining, leaves were bleached with 80% EtOH at 80°C for 10-20 min until the chlorophyll was completely removed from the leaves. The bleached leaves were stained by iodine solution containing 0.7% KI (W/V) and 0.3% I (W/V) for three minutes. The leaves were rinsed by water and photographed.

For weight measurements, several discs were obtained in replicates from ten leaves. Leaf fresh weight and leaf area were measured, then the leaves were dried at 70°C for 48 h to measure the dry weight.

### Gas exchange and chlorophyll fluorescence measurement

The net CO_2_ assimilation (A_n_), stomatal conductance (g_s_), and internal CO_2_ concentration (Ci) were measured in the leaves using a portable CO_2_ infrared gas analyzer (LI-COR-6400XT, LI-COR Environmental, USA), equipped with a chamber that controlled irradiance (1000 µmol photons m^-2^ s^-1^), temperature (20 °C), CO_2_ concentration (400 µmol mol^-1^), and flow rate (250 cm^3^ min^-1^). The measurements were tested at zheitgeber (ZT) 3, 6, and 9 h after the light on. The differences between the WT and GI-RNAi leaves were reproducible at the different ZT. Therefore, the data of ZT9 are presented here.

For chlorophyll fluorescence imaging of the leaves, we used an SPEEDZEN imaging system, and recorded an induction curve (after 30 mins of dark incubation under ambient O_2_ and CO_2_ conditions) with 2600 µmol actinic light for 3.5 minutes with 6000 µmol saturating pulse at every 30 second intervals to attain maximum NPQ levels without causing artificial damage during measurements, followed by 3.5 min of dark recovery; this experimental protocol was found to be adequate for the aspen leaves, as fluorescence had saturated after the 7^th^ flash, and had returned to low levels after the 14^th^. NPQ formed by short exposure with high light is normally known as the fast NPQ (pH dependent) whereas the residual fluorescence after dark recovery is mostly known as slow component of NPQ (Zeaxanthin/qI).

### RNA extraction and qPCR

Mature poplar leaves were harvested at zeitgeber 9 or 17, immediately frozen in liquid N_2_ and ground to a fine powder with a mortar and pestle. 100 mg powder were used for RNA extraction with CTAB extraction buffer (Chang *et al*., 1993; 2% CTAB, 100 mM Tris-HCl (pH 8.0), 25 mM EDTA, 2M NaCl, 2% PVP). The samples were incubated at 65°C for 2 min and extracted twice with an equal volume of chloroform-isoamylalcohol (24:1). Nucleic acids were precipitated at -20°C for 3 hours with ¼ volumes 10 M LiCl. Precipitate was collected by centrifugation (13000 rpm, 4°C, 20 min) and purified and DNase treated with RNeasy kit according to the manufacturer’s instructions (Qiagen). RNA integrity was confirmed by agarose gel. 1000 ng RNA were used for cDNA synthesis with iScript™ cDNA Synthesis Kit (Biorad). The cDNA was diluted 50 times for downstream applications. Quantitative real-time PCR (qPCR) was run on a LightCycler® 480 with SYBR Green I Master (Roche). All kits and machines were used according to the manufacturer’s instructions. The reaction protocol started with 5 min pre-incubation at 95°C, followed by 50 cycles of amplification consisting of 10 s denaturation at 95°C, 15 s annealing at 60°C and 20 s elongation at 72°C. For the acquisition of a melting curve fluorescence was measured during the step-wise increase in temperature from 65°C to 97°C. Relative expression levels were obtained using the 2^-ΔΔCq method (Livak & Schmittgen, 2001). GeNorm identified UBQ and 18S as most stable reference genes. All used primers had an efficiency of >1,8 and their correct product was confirmed by sequencing.

### C/N ratio measurement

Dried leaves were pooled and ground with a mortar and pestle. Dry mass was defined by oven drying at 70 °C for 24 hours. C and N concentrations (mass based) of the samples (5 mg dry weight) were determined with Elemental analyzer (Flash EA 2000, Thermo Fisher Scientific, Bremen, Germany) in four biological replicates. C and N of the dried sample material was converted to CO_2_ and N_2_ by combustion. The results were corrected for drift and sample size effect (non-linearity). Working standards were wheat and maize flours calibrated against reference standards. For ωN, atropine, cellulose, and NIST 1515 apple leaves. For ωC, cyclohexanone, nicotinamide, and sucrose.

### Outdoor experiment

For FT, the hybrid aspen plants (T89), FT-RNAi, and FT1ox lines that were characterized in Böhlenius et al. (2006), were obtained from the tissue culture and were potted in three litres of soil. The plants were kept in greenhouse subjected to 18 h photoperiod of 200 µmole light intensity. The temperature was 20/15°C (day/night). Fifty days after recovery, the plants were transferred to natural conditions outside of the greenhouse (on August 22, 2019).

For GI, two groups of trees were used in this experiment. The first group is the trees that were simply grafted on WT rootstock as described in Fig. 1B. The second group was grafted using Y method grafting as described in Fig.1B. The trees passed the first growth cycle and after the dormancy break the WT buds were removed from the first group of trees. All trees were repotted in 10 litre pots and flushed in the greenhouse in June 2020. On the beginning of July, the trees were moved out of greenhouse (Umeå, Sweden) to follow the autumnal senescence, exposed to natural conditions such as light and temperature.

### Field experiment

This experiment was a part of a larger trial of genetically modified (GMO) aspens. All genotypes were obtained from the tissue culture facility (Umeå Plant Science Centre) and potted in the greenhouse. The trees were the potted in at field site at Våxtorp, southern Sweden, in 2014. Height and chlorophyll content were measured for two year old trees.

### Statistical analyses

The multiple ways analysis of variance (ANOVA) analysis was performed using the Info-Stat/Student program (http://www.infostat.com.ar/index.php?mod=page&id=37) and the Fisher’s Least Significant Difference (LSD) were calculated for statistical analyses. As well as t-tests were used for two-group statistical analyses. A difference at P <0.05 was considered as significant.

## Acknowledgements

We thank Ulf Johansson at the Tönnersjöheden Experimental Forest, Swedish University of Agricultural Sciences, for assistance with the field experiments in southern Sweden, and Kathryn Robinson for some analyses and comments on the manuscript.

